# A dual-purpose real-time indicator and transcriptional integrator for calcium detection in living cells

**DOI:** 10.1101/2021.12.22.473920

**Authors:** Elbegduuren Erdenee, Alice Y. Ting

## Abstract

Calcium is a ubiquitous second messenger in eukaryotes, correlated with neuronal activity and T-cell activation among other processes. Real-time calcium indicators such as GCaMP have recently been complemented by newer calcium integrators that convert transient calcium activity into stable gene expression. Here we introduce LuCID, a dual-purpose real-time calcium indicator and transcriptional calcium integrator that combines the benefits of both calcium detection technologies. We show that the calcium-dependent split luciferase component of LuCID provides real-time bioluminescence readout of calcium dynamics in cells, while the GI/FKF1 split GAL4 component of LuCID converts calcium-generated bioluminescence into stable gene expression. We also show that LuCID’s modular design enables it to also read out other cellular events such as protein-protein interactions. LuCID adds to the arsenal of tools for studying cells and cell populations that utilize calcium for signaling.

## Main text

Calcium is a central player in a vast array of signaling processes, from neuron and T-cell activation^1^ to mitochondria-ER coupling^2^ and transcriptional regulation^3^. Consequently, real-time indicators for visualizing the spatiotemporal patterns of calcium signaling in living cells have been transformative for cell biology, neuroscience, immunology, and other fields^4-5^. Recently, these indicators have been complemented by a new class of calcium probes – calcium *integrators* – which convert transient calcium activity into stable signals that can be read out by large-scale microscopy, RNA-sequencing, or, if desired, altered cellular behavior. Such integrators include the calcium- and UV-light dependent photoswitchable fluorescent protein CaMPARI^6–8^ and the calcium- and blue light-gated transcription factors FLARE^9^, FLiCRE^10^, scFLARE^11^, and Cal-Light^12^.

Calcium indicators and calcium integrators each have unique and complementary strengths: indicators provide real-time, dynamic read-out over long experimental time windows with millisecond temporal resolution and high subcellular spatial resolution. Integrators record the cumulative calcium activity of cells over a single user-defined time window, but they are also highly scalable and versatile: the calcium record can be read out by microscopy, FACS, or single-cell RNA sequencing, and can also be used to drive the expression of functional proteins such as channelrhodopsin for re-activation of previously active cell subpopulations^10^. Given the complementary strengths of calcium indicators and integrators, we sought to design a dual-purpose probe that can function in either capacity, giving experimentalists maximal flexibility. In addition, we simplified our probe by removing the requirement for exogenous light. Instead, it is gated by a bio-orthogonal and nontoxic small molecule, which makes the tool potentially applicable to large and non-transparent tissue regions.

Our new probe, named LuCID, for Luciferin- and Calcium-Induced Dimerization, is a calcium and drug-gated transcription factor that gives bioluminescence readout of calcium activity as it is occurring in real-time in living cells, as well as a stable transcriptional record of that past calcium activity 12-18 hours later.

## Results and Discussion

The design of LuCID is shown in **Figure 1A**. Calcium is detected via reconstitution of a split luciferase, NanoBIT^13^, giving real-time bioluminescence readout of intracellular calcium levels. To convert NanoBiT bioluminescence to gene expression, we envisioned BRET from NanoBiT driving the activation of a light-regulated transcription factor. While our previous work showed that BRET between the luciferase NanoLuc and the photosensory domain LOV is possible^14^, it was unknown if luciferase-generated bioluminescence would be sufficient to drive the activity of existing light-dependent transcription factors.

**Figure 1.**
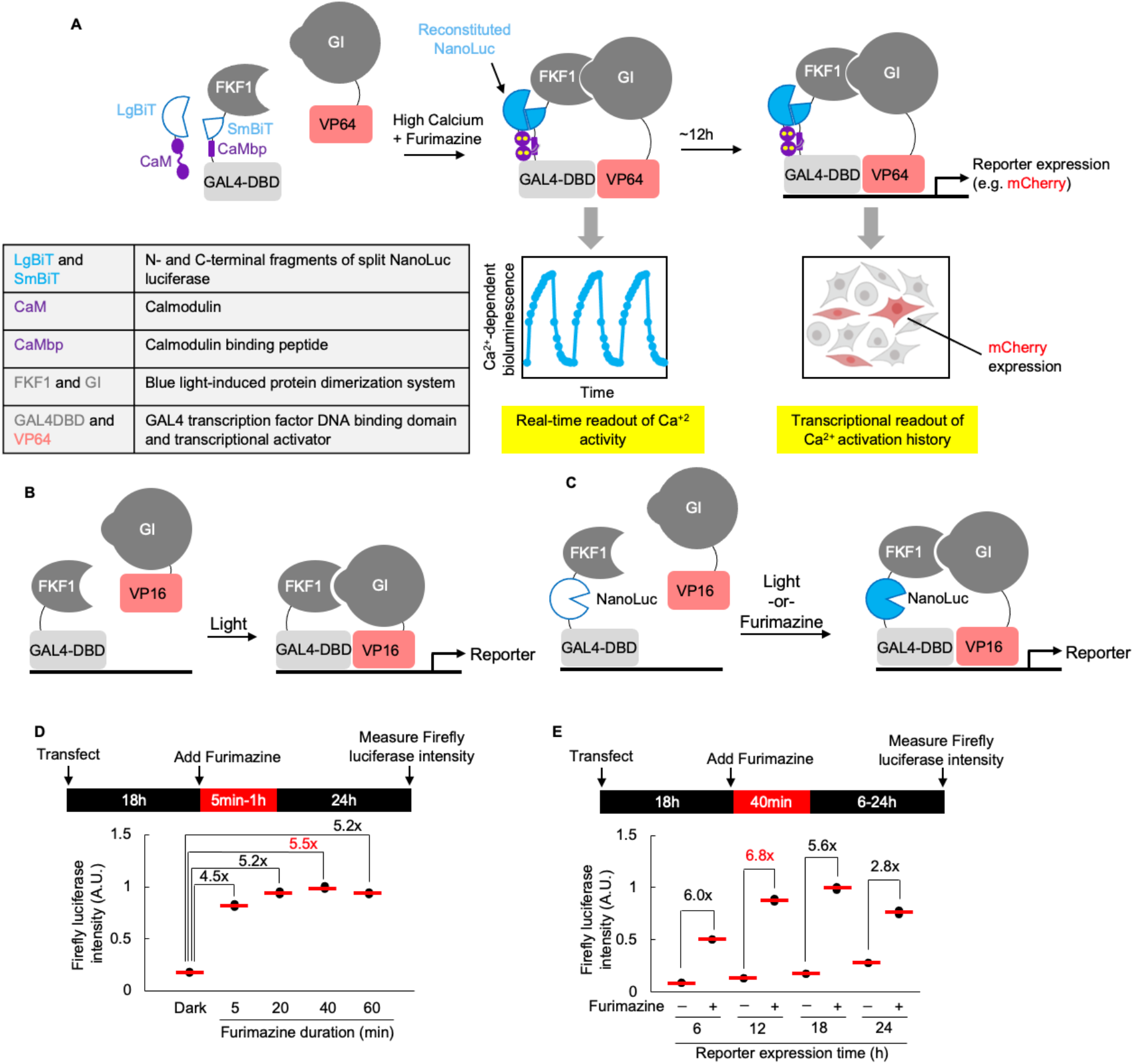
Schematic of LuCID and introduction of luciferin-gated gene expression. **A**). LuCID is a dual calcium indicator and integrator. Furimazine is the substrate used by NanoLuc to generate bioluminescence. **B)**. Light-gated split GAL4 system from Quejada et al., 2017 **C)**. Introduction of NanoLuc to constructs in (B) confers furimazine-gating. Bioluminescence from NanoLuc activates FKF1 via BRET, which recruits GI-VP16 to GAL4-DBD to drive reporter gene expression. **D)**. Test of constructs in (C) using firefly luciferase (FLuc) as the reporter gene. Furimazine was added to cells for 5 min – 1 hour, and FLuc activity was recorded 24 hours later. Two replicates per condition. Red line, mean. **E)**. Same as (D) but optimizing FLuc reporter expression times post-furimazine stimulation. Two replicates per condition. Red line, mean.

To test this, we first selected NanoLuc as our BRET donor, because it is the brightest of all known luciferases, giving 150-fold more luminescence than either Firefly or Renilla luciferases in the presence of its small-molecule luciferin substrate, furimazine^15^. Furthermore, NanoLuc does not require ATP for light generation, and it has been shown to work in a variety of *in vivo* settings including the mouse brain^16^ and liver^17^, worms^18^, and zebrafish^19^. Because NanoLuc emits blue light with a peak of ∼450 nm, it is well matched to the activation spectrum of cryptochromes that use flavin as a cofactor. Several cryptochrome-based transcription factors have been reported, including LightOn^20^, LITEZ^21^, TULIP^22^, TAEL^23^, and GI/FKF1^24,25^. We selected the GI/FKF1 system for use in LuCID because its reconstitution upon light stimulation is slow to reverse (>2hours^26^), a desirable feature for an integrator (**Figure 1B**).

To test NanoLuc regulation of GI/FKF1, we prepared genetic fusions of NanoLuc to FKF1, the light-sensitive, flavin-containing component of the GI/FKF1 system (**Figures 1C, S1A**). We expressed the components in HEK cells, along with a UAS-firefly luciferase reporter (FLuc), which is orthogonal to NanoLuc, does not use furimazine, and emits 560 nm light instead of 460 nm light. Cells were treated with furimazine for 1 hour, then cultured for 24 hours to allow time for FLuc transcription and translation. Figure S1a shows that FLuc activity was detected in cells either treated with furimazine or exposed to light, but not in untreated cells left in the dark. Similarly, expression of a different reporter gene, mCherry, was 6-fold greater in HEK293T cells treated with furimazine compared to untreated cells (**Figure S1B, C**). Fusion of NanoLuc to the N-terminus of FKF1 gave stronger turn-on than fusion to FKF1’s C-terminus (**Figure S1A**).

Optimization showed the strongest signal-to-noise ratios with a furimazine treatment time of 40 minutes and FLuc reporter expression time of 12 hours (**Figure 1D-E**). We performed a control in which NanoLuc was co-expressed with, but not directly fused to, GI/FKF1; no FLuc reporter expression was observed (**Figures S1E**). This is consistent with our previous observation that NanoLuc-LOV BRET is strongly proximity-dependent^14^, and inconsistent with a previous report^27^ showing BRET-based activation of GI/FKF1 via non-fused NanoLuc (**Figure S1D**). In **Figure S1E**, we are unable to observe furimazine-driven gene expression using this non-fused NanoLuc-GI/FKF1 system (BEACON)^27^ despite high bioluminescence from NanoLuc (**Figure S1F**) and correct nuclear localization of components (**Figures S1G**).

Next, we designed the calcium sensing component of LuCID. Previous studies have described a variety of biosensors that detect calcium with bioluminescence readout^28–30^. We tested three single-component calcium biosensors: GeNL(Ca^2+)^ based on CaM/M13 insertions into an enhanced Nano-lantern^31^, CaMBI based on CaM/M13 insertion into Antares^17^, and an unpublished circularly permuted NanoLuc (personal communication^32^). In all cases, the dynamic ranges were insufficient, primarily due to high background in the absence of calcium (**Figure S2A-B**). We hypothesized that a two-component sensor in which calcium drives the reconstitution of a split luciferase, as in Nguyen et al., 2020^33^, may have greater dynamic range due to its requirement for an intermolecular rather than an intramolecular interaction.

We utilized the split NanoLuc system called NanoBiT, in which the two split fragments, LgBiT (18 kDa, amino acids 1-156) and SmBiT (1.3 kDa, amino acids 157-167) have low affinity for one another (K_d_ = 190 µM) and must be driven together to enable reconstitution of luciferase activity^13^. We fused CaM to LgBiT and M13 to SmBiT and measured the bioluminescence of reconstituted NanoLuc in HEK293T cells treated with CaCl_2_ and ionomycin or left untreated. We observed a +/- Ca^2+^ signal ratio of 48, in contrast to the single component calcium biosensors which gave a maximum +/- Ca^2+^ signal ratio of 1.4 (**Figure S2B**).

Encouraged by this large dynamic range, we inserted the NanoBiT fusions into the GI/FKF1 system. We generated 18 distinct construct combinations (D1-18) with varying geometries (**Figures 2B, S2C**) guided by the following general principles: (1) LgBiT or SmBiT were placed close to FKF1 to enable efficient BRET, (2) the GI-VP16 component was held constant, (3) smaller protein domains were sandwiched between GAL4DBD and FKF1 in order to reduce the distance between them and maintain ability of GI/FKF1 interaction to drive GAL4DBD-VP16 reconstitution, and (4) we avoided fusions to the C-terminus of FKF1, based on our data with full-length NanoLuc (**Figure S1A**).

**Figure 2.**
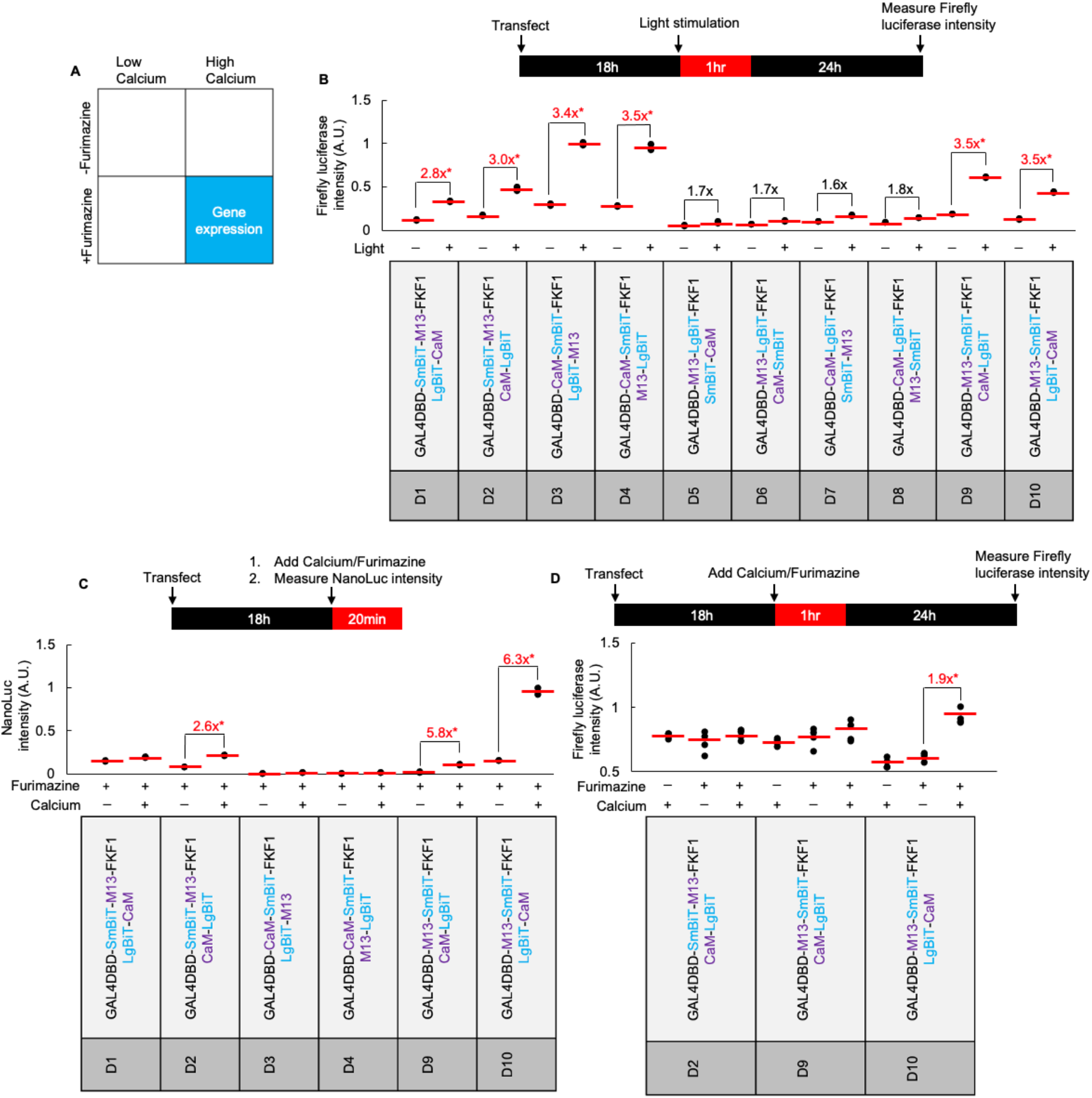
Introducing calcium gating. **A)**. LuCID is an AND logic gate, requiring both calcium and furimazine to produce gene expression. **B)**. Optimizing LuCID geometry. Construct pairs D1-D10 were tested in HEK293T cells with 1 hour of light stimulation. Eight additional pairs (D11-18) tested in **Figure S2C**. *best pairs. Two replicates per condition. Red line, mean. **C)**. The 6 best pairs from (B) were tested with 20 minutes of furimazine +/- calcium stimulation (2µM ionomycin and 5mM CaCl2) in HEK293T cells. NanoLuc bioluminescence was recorded ∼4-6min after calcium and furimazine addition. Two replicates per condition. **D)**. FLuc reporter readout 24 hours post-stimulation, for three best pairs (starred) from (C). Four replicates per condition. Red line, mean.

All 18 construct combinations were first tested for their ability to drive light-dependent gene expression (**Figures 2B, S2C**). The six pairs that gave 2.5-fold or greater +/- light FLuc reporter activity were then tested for calcium-dependent bioluminescence (**Figure 2C**). The three best pairs had either C-terminal or N-terminal exposed CaM domains, which may facilitate intermolecular complexation with M13 fusions. These three pairs were then expressed in HEK293T cells along with UAS-FLuc reporter (**Figure 2D**). The pair with the best +/- calcium FLuc expression was further optimized in terms of calcium/furimazine treatment times (**Figure 3A**) and concentrations (**Figure S3C-F**), reporter expression times (**Figure 3B**), and plasmid ratios (**Figure 3C**). Finally, we replaced the VP16 transcriptional activator domain with VP64, a stronger activator, to produce LuCID. Under optimized conditions, LuCID’s +/- Ca^2+^ signal ratio was ∼16.5 in HEK293T cells (**Figure 3C**).

**Figure 3.**
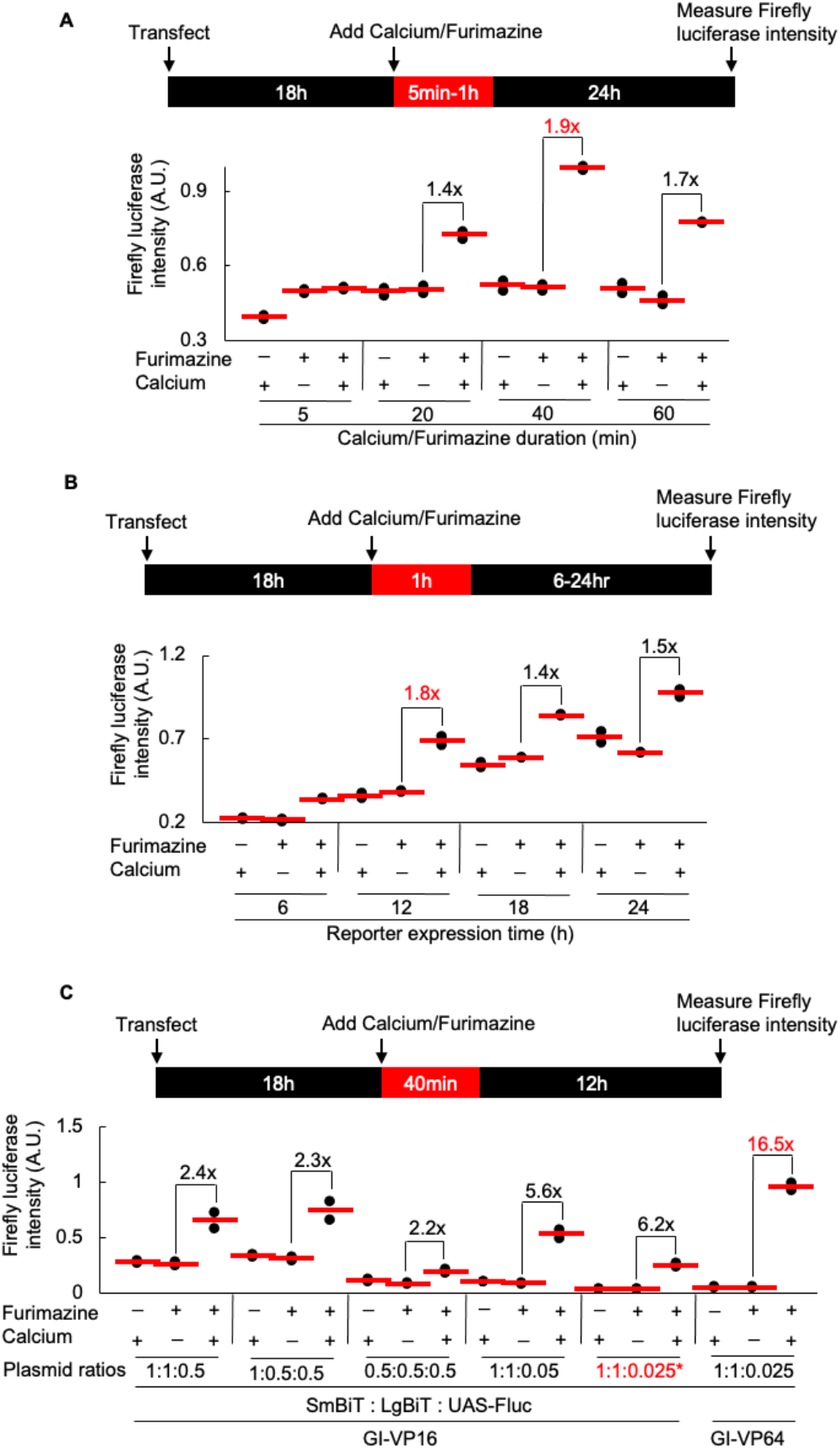
Optimization of LuCID dynamic range. **A)** Optimizing stimulation time. FLuc expression in HEK293T cells measured 24 hours post-stimulation. Two replicates per condition. **B)** Optimizing FLuc reporter expression time. Two replicates per condition. **C)** Testing different plasmid ratios and transcriptional activator domains (VP16 vs. VP64). SmBiT = GI-VP16-IRES-GAL4DBD-M13-SmBiT-FKF1. LgBiT = LgBiT-CaM. Stimulation time 40 minutes and FLuc activity measured 12 hours later. Two replicates per condition.

LuCID’s design enables it to read out real time calcium dynamics in addition to integrating calcium spikes into a stable transcriptional signal, because LuCID’s calcium sensor – the calcium-regulated split NanoLuc – gives an instantaneous bioluminescence readout and is fully and rapidly reversible. To test LuCID’s function as a real-time Ca^2+^ indicator, we simulated Ca^2+^ spikes in HEK293T cells with ionomycin + CaCl_2_ and measured LuCID bioluminescence concurrently with the fluorescence of a “gold standard Ca^2+^ indicator”, GcaMP6s^34^. We found that both readouts were well-matched (**Figure 4A**). Interestingly, the on-rate kinetics of LuCID were slower compared to GcaMP6s, while the off-rate kinetics were faster. This is likely because LuCID’s design is intermolecular while GcaMP6s is intramolecular.

**Figure 4.**
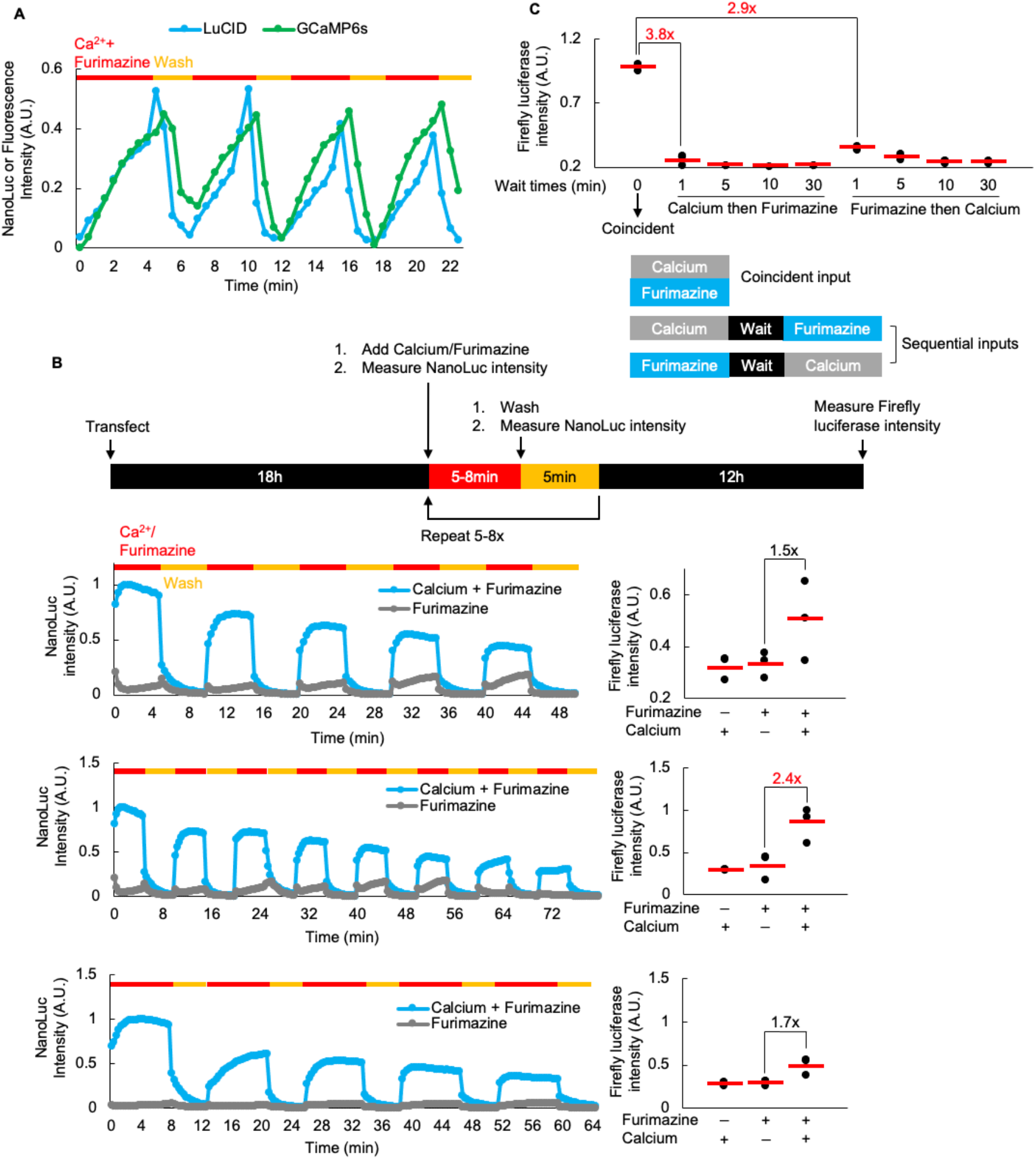
LuCID for measurement of real-time calcium dynamics. **A)**. Simultaneous measurement by LuCID (blue) and GCaMP6s (green) expressed together in HEK293T cells. Cells were stimulated repeatedly with 2µM ionomycin, 10mM CaCl2, and 15µM furimazine for 4 minutes (red bars) and washed for 2 min (yellow bars), while bioluminescence was recorded at 460 nm for LuCID and fluorescence was recorded at 512 nm (497 nm excitation) for GcaMP6s. This experiment was repeated three times. One representative trace shown. **B)** Demonstration of the dual functionality of LuCID. Matched samples were stimulated repeatedly with calcium and furimazine as indicated. In each set, one sample was measured in real-time (time traces at left) while the other sample was measured for FLuc reporter expression 12h later (graphs at right). Three replicates per condition. **C)**. LuCID requires coincident activation by calcium and furimazine. HEK293T cells expressing LuCID were stimulated sequentially with calcium (20 minutes) followed by furimazine (20 minutes) or the reverse. Intervening wait times ranged from 1-30 minutes. Three replicates per condition. Red line, mean.

We also tested whether LuCID could detect non-continuous calcium spikes in HEK293T cells. We stimulated HEK293T cells with three different “spike” patterns (**Figure 4B**). We measured both real-time calcium activity from each of these spike patterns (**Figure 4B, left**) and calcium integration 12 hours later by checking for UAS-FLuc reporter expression (**Figure 4B, right**). S/N in reporter expression is much lower for these spike-pattern treatments (highest S/N of 2.4 with 8 spikes of 5 minutes of calcium/furimazine) compared to the continuous calcium/furimazine treatment condition (**Figure 3C**; S/N = 16.5), which suggests that the M13-SmBiT/LgBiT-CaM component of LuCID is so quickly reversible after washout of calcium/furimazine that the FKF1/G1 cannot be sustainably engaged. This is consistent with previous research showing that the interaction between FKF1 and GI has a slow on-rate, on the timescale of minutes^24^.

We further characterized LuCID’s performance as a calcium integrator. If LuCID functions as an “AND” logic gate, or coincidence detector, as we have designed, then the staggering of calcium and furimazine inputs should not produce reporter gene expression. **Figure 4C** shows that when furimazine precedes calcium by 1 minute, or vice versa, LuCID’s resulting signal output is significantly reduced. LuCID is a coincidence detector of calcium and furimazine due to the reversibility of its calcium sensing and the speed of furimazine washout from cells.

To highlight LuCID’s versatility in terms of readout, we also used a UAS-mCherry reporter and microscopy to detect LuCID turn on. 40 minutes of calcium and furimazine treatment induced significant mCherry reporter expression in HEK293T cells ∼18 hours later (**Figure S3A-B**). Cells receiving either calcium only or furimazine only treatment exhibited minimal mCherry expression. The mean mCherry-to-HA fluorescence intensity ratio (mCherry/HA) for the double-positive +calcium/+furimazine condition was 12.7-fold higher than the single-positive calcium-only or furimazine-only conditions. An HA tag fused to FKF1 allowed us to confirm equal expression levels across conditions using antibody staining (**Figure S3A**).

To test if LuCID could detect intracellular calcium changes within physiological ranges, we treated HEK293T cells with a Sarcoplasmic/Endoplasmic Reticulum Calcium ATPase (SERCA) activator, CDN1163 (CDN), to reduce intracellular calcium, followed by one of three well-studied SERCA inhibitors (cyclopiazonic acid (CPA), 2,5-di-(tert-butyl)-1,4-hydroquinone (DBHQ) or thapsigargin (Thaps.)) to release cytosolic calcium from internal calcium stores (**Figure 5A-B**). These inhibitors by themselves resulted in 3.6-, 4.6-, and 4.6-fold LuCID activation, respectively, for CPA, DBHQ and thapsigargin compared to furimazine-only treatment (**Figure 5A**). CDN pre-treatment reduced LuCID activation by 1.6-fold when used with CPA, 2.5-fold when used with DBHQ, and 2.6-fold when used with thapsigargin (**Figure 5A**). We are also able to measure the real-time effects of these combinations of drug treatments, where we see significant reductions in LuCID’s real-time bioluminescence signals when cells are pre-treated with CDN compared to SERCA inhibitors alone (**Figure 5B**).

**Figure 5.**
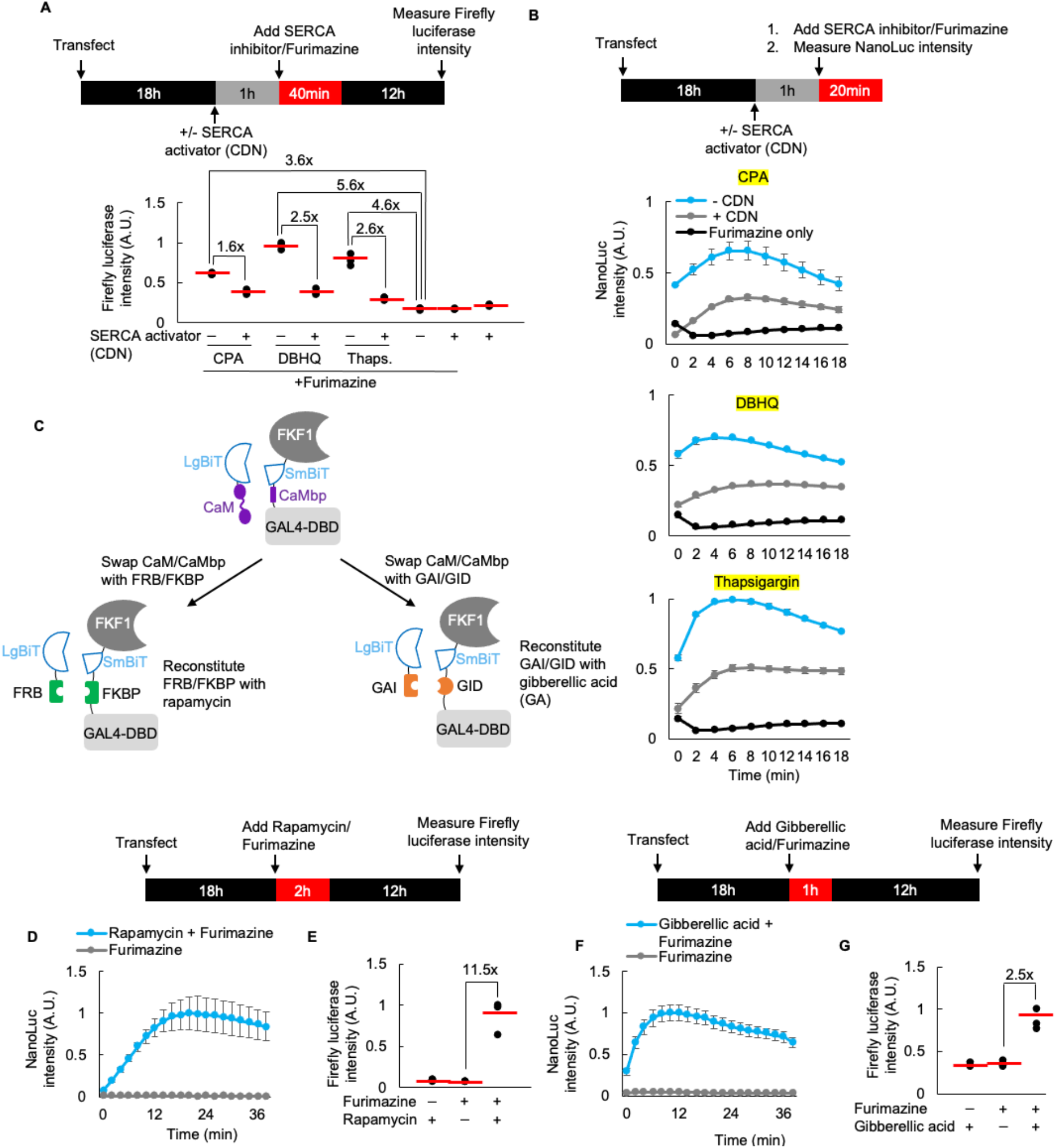
LuCID for detection of physiological calcium and protein-protein interactions. **A**). HEK293T cells expressing LuCID were treated with the SERCA activator CDN1163 (CDN) for 1 hour, then one of three SERCA inhibitors (CPA (cyclopiazonic acid), DBHQ (2, 5-di(tert-butyl) hydroquinone), or thapsigargin) for 40 minutes together with furimazine. FLuc expression was measured 12 hours later. Three replicates per condition. Red line, mean. **B)**. Real-time calcium measurements from cells treated as in (A). Error bars = +/- 1 SD **C)**. Schematic showing replacement of calcium-responsive domains of LuCID with protein-protein interaction partners. **D)**. Real-time bioluminescence measurements of rapamycin-triggered FRB-FKBP interactions with LuCID in HEK293T cells. Three replicates per condition. Error bars = +/- 1 SD. **E)**. Same as D, but FLuc reporter expression was measured 12 hours later. Three replicates per condition. Red line, mean. **F)**. Real-time bioluminescence measurements of gibberellic acid-triggered GID/GAI interactions with LuCID in HEK293T cells. Three replicates per condition. Error bars = +/- 1 SD. **G)**. Same as F, but FLuc activity was measured 12 hours later. Three replicates per condition. Red line, mean.

Finally, due to the modularity of LuCID’s design, we tested if it could be adapted for the detection of other biochemical events besides Ca^2+^ rises, such as protein-protein interactions (PPIs). To test this, we replaced the CaM and CaMbp calcium-sensing domains of LuCID with the chemically-induced protein dimerizing domains FRB and FKBP^35^ or GAI and GID^36^ (**Figure 5C**). **Figure 5E** shows that 2 hours of rapamycin and furimazine treatment of HEK293T cells expressing the FKBP/FRB reporter induces 11.5-fold higher FLuc expression than rapamycin-only or furimazine-only treatments. 2 hours of gibberellic acid (GA) and furimazine treatment of the GAI/GID set induces 2.5-fold higher FLuc expression than GA-only or furimazine-only treatments (**Figure 5G**). As with calcium-sensing LuCID, real-time PPI dynamics of both the rapamycin- and gibberellic-acid-based systems could also be measured (**Figure 5D, F**).

In summary, LuCID is a dual-purpose calcium probe that can be used both for real-time bioluminescence readout of calcium dynamics and stable recording of past calcium activity in the form of new gene expression. We perform extensive protein engineering and optimization to develop LUCID, characterize its performance in cell culture, use it in a small-scale screen of SERCA inhibitors, and show that the design can be extended to the detection of other cellular events such as protein-protein interactions.

Compared to other calcium integrators, such as FLiCRE developed by our lab^9–11^, and others^6–8,12^ LuCID has the advantage of being temporally gated by a small-molecule (furimazine) rather than external light. This makes it potentially easier to use in vivo, across large swaths of tissue, and also for high-throughput robotics-assisted screens. Previous work^16,17^ has shown that furimazine can access the mouse brain and liver to produce bioluminescence. Compared to real-time calcium indicators, LuCID’s main advantages are the lack of requirement for external excitation light, lack of photobleaching, and considerably higher dynamic range than single-chain calcium bioluminescence indicators such as CaMBI^17^.

LuCID does have important limitations however. Compared to FLiCRE, for example, LuCID has poorer temporal resolution, requiring 40 minutes of stimulation compared to 1 minute^10^. This is partly because furimazine delivery/washout are slower than light delivery, but we believe the primary reason is that FKF1-GI requires several minutes of sustained NanoLuc bioluminescence to become activated. Improved transcriptional activators and improved geometry that optimizes BRET between NanoLuc and transcription factor may improve temporal resolution in future designs. A second limitation is that LuCID has multiple components, one of which is large (GI is ∼129 kDa) and difficult to package even into a lentiviral vector. The tool could benefit from simplification, as we did to produce single-chain FLARE^11^, so that it is more robust and exhibits less expression level and plasmid ratio variation across experiments. Despite these challenges, LuCID does offer new capabilities among the constellation of calcium detection technologies and provides a new way to measure and study calcium signaling in biology.

## Methods

### Stimulation of HEK293T cells with light, furimazine, and/or calcium

HEK293T cells cultured and transfected as described in Supplementary Methods were stimulated as follows. For experiments with full-length NanoLuc (**Figures 1 and S1**) and BEACON^27^ (**Figure S1**), media in each well was replaced with 100µL (for 96-well plate format), 200µL (for 48-well plate format) or 400µL (for 24-well plate format) complete DMEM with or without 10µM furimazine (Promega). Cells were incubated at 37°C under 5% CO_2_ either in the dark, wrapped in aluminum foil, or stimulated with 467nm blue light at 60mW/cm^2^ and 50% duty cycle (3s of light every 6s) by placing the entire plate directly above an LED light array for the indicated times. For experiments with calcium stimulation (**Figures 2 and S2**), media in each well was replaced with 100µL (for 96-well plate format) or 200µL (for 48-well plate format) complete DMEM in addition to 2µM ionomycin (Sigma-Aldrich) and 5mM CaCl_2_ with or without 10µM furimazine (unless indicated otherwise). After stimulation, media was removed from the wells and replaced with the same volume of complete DMEM and the cells were returned to the 37°C incubator wrapped in aluminum foil for reporter expression. All manipulations were done under red light. For pulsed stimulation experiment in Figure 4, 100µL complete DMEM containing 2µM ionomycin and 10mM CaCl_2_ with or without 15µM furimazine was added to each well for the indicated times and then replaced with complete DMEM without additional drugs during the indicated washes. Calcium/furimazine addition and wash steps were repeated 4-8 times to simulate calcium spikes.

### Real-time bioluminescence measurements with LuCID and NanoLuc

Real-time bioluminescence measurements from either full-length NanoLuc or reconstituted NanoBiT in LuCID were taken on a Tecan Infinite M1000 Pro platereader immediately after adding complete DMEM with 2µM ionomycin + 5mM CaCl_2_ with or without 10µM furimazine (unless otherwise indicated) to each well. Measurements were taken using the blue filter (450nm) with 1000ms integration time every 2 minutes for 20 minutes. Bioluminescence from the pulsed experiment in Figure 4 was measured both immediately after adding complete DMEM with 2µM ionomycin + 10mM CaCl_2_ with or without 15µM furimazine and also after adding complete DMEM without additional drugs using the platereader’s kinetic cycle function with the following settings: 10 cycles, 500ms kinetic interval with 100ms integration time with 450nm blue filter. For real-time calcium measurements with GCaMP6s, fluorescence signal was measured immediately after adding complete DMEM with 2µM ionomycin + 10mM CaCl_2_ with or without 15µM furimazine when GCaMP6s was co-expressed with LuCID (**Figure 4A**) or after adding complete DMEM with 2µM ionomycin + 10mM CaCl_2_ when GCaMP6s was expressed alone (**Figure S4**) using the same kinetic cycle as above. Measurements were recorded with 497nm excitation and 512nm emission wavelengths.

### Measurement of LuCID-driven gene expression

12-24 hours after calcium/furimazine stimulation as described above, firefly luciferase (FLuc) reporter gene expression was measured by luminescence on a platereader (Tecan Infinite M1000 Pro) using the Bright-Glo Luciferase Assay System (Promega). The Bright-Glo reagent was thawed at room temperature and diluted two-fold with PBS. Media was aspirated from each well and 50µL of the diluted Bright-Glo reagent was added to each well. Luminescence was immediately analyzed at 25°C using a 1000 ms integration time, the Green-1 filter (520-570nm) and linear shaking for 10 s.

### SERCA inhibitor/activator experiments (Figure 5)

HEK293T cells were transfected as indicated in Table 2. After 18-hour incubation wrapped in foil in a 37°C/5% CO_2_ incubator, media was replaced with 100µL complete DMEM +/- 30µM SERCA activator CDN1163 (Sigma-Aldrich) and incubated at 37°C for 1 hour. Then, media was replaced again with 100µL DMEM + 10% FBS + 1% PS containing 10µM furimazine and one of the following SERCA inhibitors: 10µM cyclopiazonic acid (CPA) (Sigma-Aldrich), 10µM 2,5-di-(tert-butyl)-1,4-hydroquinone (DBHQ) (Sigma-Aldrich), or 10µM Thapsigargin (EMD Millipore). Bioluminescence measurements were taken in real-time immediately after adding SERCA inhibitor + furimazine and compared to cells treated with just 10µM furimazine. Firefly luciferase reporter expression was measured ∼12 hours later, as described above.

### Rapamycin and gibberellic acid experiments (Figure 5)

HEK293T cells were transfected as indicated in Table 2. After 18-hour incubation, media in each well was replaced with 100µL complete DMEM with the following drug combinations: 1) 100nM rapamycin only, 2) 10µm furimazine only and 3) 100nm rapamycin + 10µM furimazine combined for 2hrs at 37°C under 5% CO_2_ or with 1) 10µM gibberellic acid (GA) only, 2) 10µm furimazine only and 3) 10µM GA + 10µM furimazine combined for 2hrs at 37°C under 5% CO_2_ prior to luminescence readout 12 hours later, respectively for each experiment. Real-time measurements were taken immediately after adding the above drug combinations on the platereader.

See Supporting Information and **Tables 1-2** for additional methods and information on cloning, cell culture and transfection, cell fixation, immunostaining, and fluorescence microscopy.

## Acknowledgements

This work was supported by the NIH (R01 MH119353 to A.Y.T.)

## References

1. Fracchia KM, Pai CY, Walsh CM. Modulation of T Cell Metabolism and Function through Calcium Signaling. Front Immunol. 2013;4. doi:10.3389/fimmu.2013.00324

2. Filadi R, Theurey P, Pizzo P. The endoplasmic reticulum-mitochondria coupling in health and disease: Molecules, functions and significance. Cell Calcium. 2017;62:1–15. doi:10.1016/j.ceca.2017.01.003

3. Hogan PG. Transcriptional regulation by calcium, calcineurin, and NFAT. Genes Dev. 2003;17(18):2205–2232. doi:10.1101/gad.1102703

4. Rose T, Goltstein PM, Portugues R, Griesbeck O. Putting a finishing touch on GECIs. Front Mol Neurosci. 2014;7. doi:10.3389/fnmol.2014.00088

5. Shen Y, Nasu Y, Shkolnikov I, Kim A, Campbell RE. Engineering genetically encoded fluorescent indicators for imaging of neuronal activity: Progress and prospects. Neurosci Res. 2020;152:3–14. doi:10.1016/j.neures.2020.01.011

6. Fosque BF, Sun Y, Dana H, et al. Labeling of active neural circuits in vivo with designed calcium integrators. Science. 2015;347(6223):755–760. doi:10.1126/science.1260922

7. Moeyaert B, Holt G, Madangopal R, et al. Improved methods for marking active neuron populations. Nat Commun. 2018;9(1):4440. doi:10.1038/s41467-018-06935-2

8. Sha F, Abdelfattah AS, Patel R, Schreiter ER. Erasable labeling of neuronal activity using a reversible calcium marker. eLife. 2020;9:e57249. doi:10.7554/eLife.57249

9. Wang W, Wildes CP, Pattarabanjird T, et al. A light- and calcium-gated transcription factor for imaging and manipulating activated neurons. Nat Biotechnol. 2017;35(9):864–871. doi:10.1038/nbt.3909

10. Kim CK, Sanchez MI, Hoerbelt P, et al. A Molecular Calcium Integrator Reveals a Striatal Cell Type Driving Aversion. Cell. 2020;183(7):2003-2019.e16. doi:10.1016/j.cell.2020.11.015

11. Sanchez MI, Nguyen QA, Wang W, Soltesz I, Ting AY. Transcriptional readout of neuronal activity via an engineered Ca 2+ -activated protease. Proc Natl Acad Sci. 2020;117(52):33186–33196. doi:10.1073/pnas.2006521117

12. Lee D, Hyun JH, Jung K, Hannan P, Kwon HB. A calcium- and light-gated switch to induce gene expression in activated neurons. Nat Biotechnol. 2017;35(9):858–863. doi:10.1038/nbt.3902

13. Dixon AS, Schwinn MK, Hall MP, et al. NanoLuc Complementation Reporter Optimized for Accurate Measurement of Protein Interactions in Cells. ACS Chem Biol. 2016;11(2):400–408. doi:10.1021/acschembio.5b00753

14. Kim CK, Cho KF, Kim MW, Ting AY. Luciferase-LOV BRET enables versatile and specific transcriptional readout of cellular protein-protein interactions. eLife. 2019;8:e43826. doi:10.7554/eLife.43826

15. Hall MP, Unch J, Binkowski BF, et al. Engineered Luciferase Reporter from a Deep Sea Shrimp Utilizing a Novel Imidazopyrazinone Substrate. ACS Chem Biol. 2012;7(11):1848–1857. doi:10.1021/cb3002478

16. Germain-Genevois C, Garandeau O, Couillaud F. Detection of Brain Tumors and Systemic Metastases Using NanoLuc and Fluc for Dual Reporter Imaging. Mol Imaging Biol. 2016;18(1):62–69. doi:10.1007/s11307-015-0864-2

17. Oh Y, Park Y, Cho JH, et al. An orange calcium-modulated bioluminescent indicator for non-invasive activity imaging. Nat Chem Biol. 2019;15(5):433–436. doi:10.1038/s41589-019-0256-z

18. Sfarcic I, Bui T, Daniels EC, Troemel ER. Nanoluciferase-Based Method for Detecting Gene Expression in Caenorhabditis elegans. Genetics. 2019;213(4):1197–1207. doi:10.1534/genetics.119.302655

19. Dumesic PA, Egan DF, Gut P, et al. An Evolutionarily Conserved uORF Regulates PGC1α and Oxidative Metabolism in Mice, Flies, and Bluefin Tuna. Cell Metab. 2019;30(1):190-200.e6. doi:10.1016/j.cmet.2019.04.013

20. Wang X, Chen X, Yang Y. Spatiotemporal control of gene expression by a light-switchable transgene system. Nat Methods. 2012;9(3):266–269. doi:10.1038/nmeth.1892

21. Polstein LR, Gersbach CA. Light-Inducible Gene Regulation with Engineered Zinc Finger Proteins. In: Cambridge S, ed. Photoswitching Proteins. Vol 1148. Methods in Molecular Biology. Springer New York; 2014:89–107. doi:10.1007/978-1-4939-0470-9_7

22. Strickland D, Lin Y, Wagner E, et al. TULIPs: tunable, light-controlled interacting protein tags for cell biology. Nat Methods. 2012;9(4):379–384. doi:10.1038/nmeth.1904

23. Reade A, Motta-Mena LB, Gardner KH, Stainier DY, Weiner OD, Woo S. TAEL: A zebrafish-optimized optogenetic gene expression system with fine spatial and temporal control. Development. Published online January 1, 2016:dev.139238. doi:10.1242/dev.139238

24. Yazawa M, Sadaghiani AM, Hsueh B, Dolmetsch RE. Induction of protein-protein interactions in live cells using light. Nat Biotechnol. 2009;27(10):941–945. doi:10.1038/nbt.1569

25. Quejada JR, Park SHE, Awari DW, et al. Optimized light-inducible transcription in mammalian cells using Flavin Kelch-repeat F-box1/GIGANTEA and CRY2/CIB1. Nucleic Acids Res. 2017;45(20):e172–e172. doi:10.1093/nar/gkx804

26. Seong J, Lin MZ. Optobiochemistry: Genetically Encoded Control of Protein Activity by Light. Annu Rev Biochem. 2021;90(1):475–501. doi:10.1146/annurev-biochem-072420-112431

27. Parag-Sharma K, O’Banion CP, Henry EC, et al. Engineered BRET-Based Biologic Light Sources Enable Spatiotemporal Control over Diverse Optogenetic Systems. ACS Synth Biol. 2020;9(1):1–9. doi:10.1021/acssynbio.9b00277

28. Baubet V, Le Mouellic H, Campbell AK, Lucas-Meunier E, Fossier P, Brulet P. Chimeric green fluorescent protein-aequorin as bioluminescent Ca2+ reporters at the single-cell level. Proc Natl Acad Sci. 2000;97(13):7260–7265. doi:10.1073/pnas.97.13.7260

29. Markova SV, Vysotski ES, Blinks JR, Burakova LP, Wang BC, Lee J. Obelin from the Bioluminescent Marine Hydroid Obelia geniculata : Cloning, Expression, and Comparison of Some Properties with Those of Other Ca 2+ -Regulated Photoproteins. Biochemistry. 2002;41(7):2227–2236. doi:10.1021/bi0117910

30. Saito K, Chang YF, Horikawa K, et al. Luminescent proteins for high-speed single-cell and whole-body imaging. Nat Commun. 2012;3(1):1262. doi:10.1038/ncomms2248

31. Suzuki K, Kimura T, Shinoda H, et al. Five colour variants of bright luminescent protein for real-time multicolour bioimaging. Nat Commun. 2016;7(1):13718. doi:10.1038/ncomms13718

32. Kim CK. Circularly-permuted NanoLuc-based calcium sensor.

33. Nguyen LP, Nguyen HT, Yong HJ, et al. Establishment of a NanoBiT-Based Cytosolic Ca 2+ Sensor by Optimizing Calmodulin-Binding Motif and Protein Expression Levels. Mol Cells. 2020;43(11):909–920. doi:10.14348/molcells.2020.0144

34. Chen TW, Wardill TJ, Sun Y, et al. Ultrasensitive fluorescent proteins for imaging neuronal activity. Nature. 2013;499(7458):295–300. doi:10.1038/nature12354

35. Rivera VM, Clackson T, Natesan S, et al. A humanized system for pharmacologic control of gene expression. Nat Med. 1996;2(9):1028–1032. doi:10.1038/nm0996-1028

36. Gao Y, Xiong X, Wong S, Charles EJ, Lim WA, Qi LS. Complex transcriptional modulation with orthogonal and inducible dCas9 regulators. Nat Methods. 2016;13(12):1043–1049. doi:10.1038/nmeth.4042

