## Supporting Information for "A dual-purpose real-time indicator and transcriptional integrator for calcium detection in living cells"

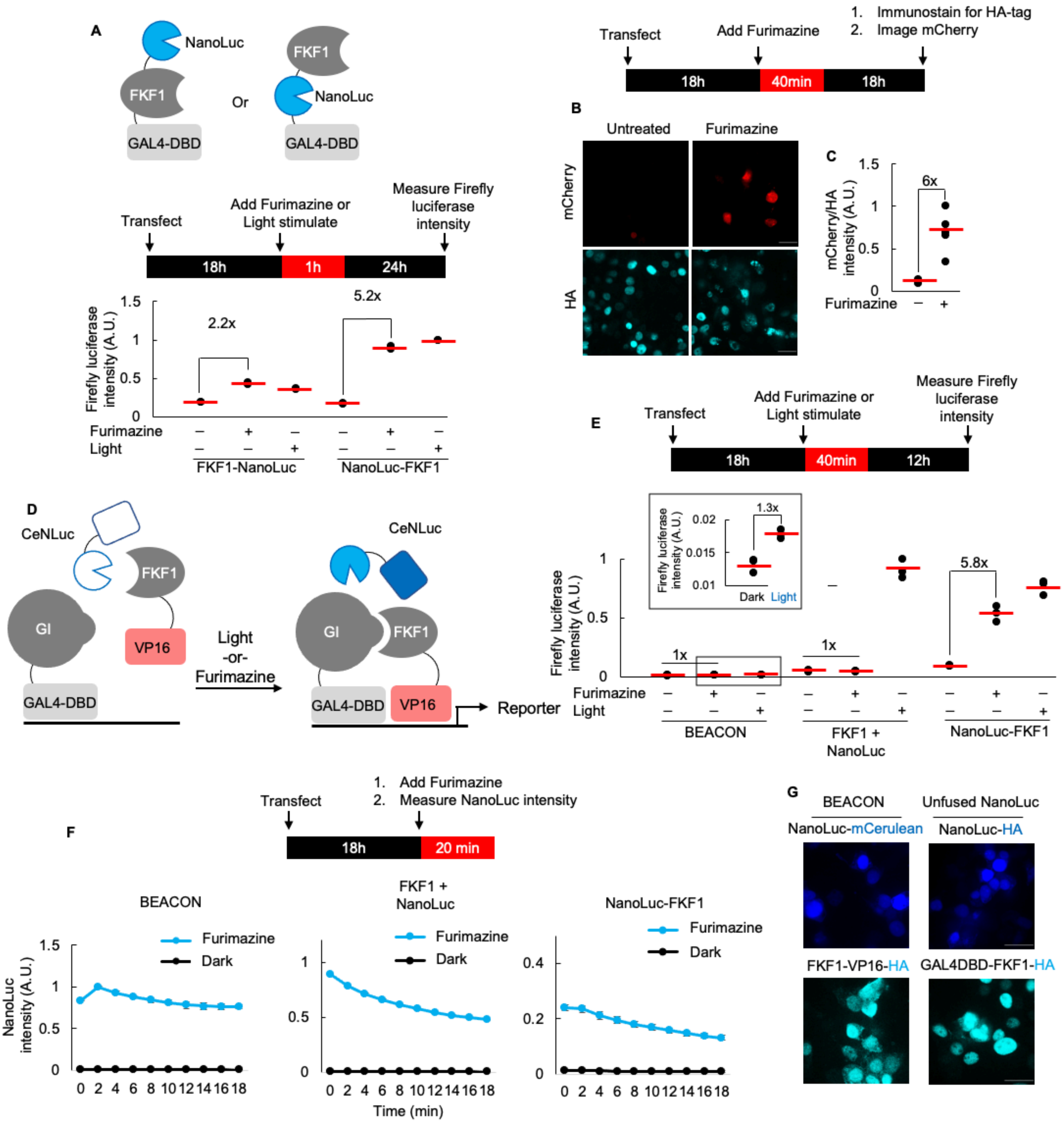
A
NanoLuc
FKF1
GAL4-DBD
Or
FKF1
NanoLuc
GAL4-DBD
Transfect
Add Furimazine or Light stimulate
Measure Firefly luciferase intensity
18h
1h
24h
Firefly luciferase intensity (A.U.)
1.5
1
0.5
0
2.2x
5.2x
Furimazine
Light
FKF1-NanoLuc
NanoLuc-FKF1
Transfect
Add Furimazine
1. Immunostain for HA-tag
2. Image mCherry
18h
40min
18h
B
Untreated
Furimazine
mCherry
HA
C
mCherry/HA intensity (A.U.)
1.5
1
0.5
0
6x
Furimazine
D
CeNLuc
FKF1
GI
VP16
GAL4-DBD
Light
-or-
Furimazine
CeNLuc
GI
FKF1
GAL4-DBD
VP16
Reporter
E
Transfect
Add Furimazine or Light stimulate
Measure Firefly luciferase intensity
18h
40min
12h
Firefly luciferase intensity (A.U.)
0.02
0.015
0.01
1.3x
Dark
Light
1
0.5
0
1x
1x
5.8x
Furimazine
Light
BEACON
FKF1 + NanoLuc
NanoLuc-FKF1
F
Transfect
1. Add Furimazine
2. Measure NanoLuc intensity
18h
20 min
BEACON
Furimazine
Dark
NanoLuc intensity (A.U.)
1.5
1
0.5
0
0 2 4 6 8 10 12 14 16 18
FKF1 + NanoLuc
Furimazine
Dark
1
0.5
0
0 2 4 6 8 10 12 14 16 18
Time (min)
NanoLuc-FKF1
Furimazine
Dark
0.4
0.2
0
0 2 4 6 8 10 12 14 16 18
G
BEACON
Unfused NanoLuc
NanoLuc-mCerulean
NanoLuc-HA
FKF1-VP16-HA
GAL4DBD-FKF1-HA

**Supplementary figure 1 (related to figure 1)**. **Testing full-length NanoLuc insertions into GI/FKF1 and comparison to BEACON A**). Two different geometries tested for full-length NanoLuc fusion to GAL4DBD-FKF1. Constructs were expressed in HEK 293T cells with firefly luciferase (FLuc) as the reporter gene. Cells were stimulated with either light or furimazine for 1 hour. Two replicates per condition. **B**). Example immunofluorescence images of mCherry expression in HEK293T cells 18 hours following furimazine treatment for 40 min (top, red). HA antibody staining labels GAL4DBD-NanoLuc-FKF1 (bottom, cyan). Scale bar = 30 μm. **C)**. Mean mCherry/HA fluorescence intensity ratio for all HA-positive HEK293T cells across 5 fields of view for experiment in (B). **D)**. Schematic of BEACON (Parag-Sharma et al., 2019). NanoLuc is fused to mCerulean3 fluorescent protein (CeNLuc). Addition of furimazine excites mCerulean3 via BRET and drives FKF1-GI interaction via FRET, reconstituting VP16-GAL4DBD. **E).** Comparison of BEACON in HEK293T cells side-by-side with GI-VP16/GAL4DBD-NanoLuc-FKF1 (right) and with GI-VP16/GAL4DBD-FKF1 with unfused NanoLuc (middle). Cells were stimulated with either light or furimazine for 40min. FLuc reporter readout was quantified 12 hours later. Inset shows zoom of boxed region; BEACON gave 1.3x increase in FLuc expression following light stimulation. Three replicates per condition. **F)**. Real-time NanoLuc intensity measurements +/- furimazine from the three conditions in (E). Trace for the average of three replicates per condition. Error bars = +/- 1 SD. **G**). Immunofluorescence images showing nuclear localization of CeNluc (NanoLuc fused to mCerulean) from BEACON (top left) and unfused HA-tagged NanoLuc (top right), HA-tagged FKF1-VP16 component from BEACON (bottom left) and HA-tagged GAL4DBD-FKF1 component used with unfused NanoLuc (bottom right). Scale bar = 30 μm.

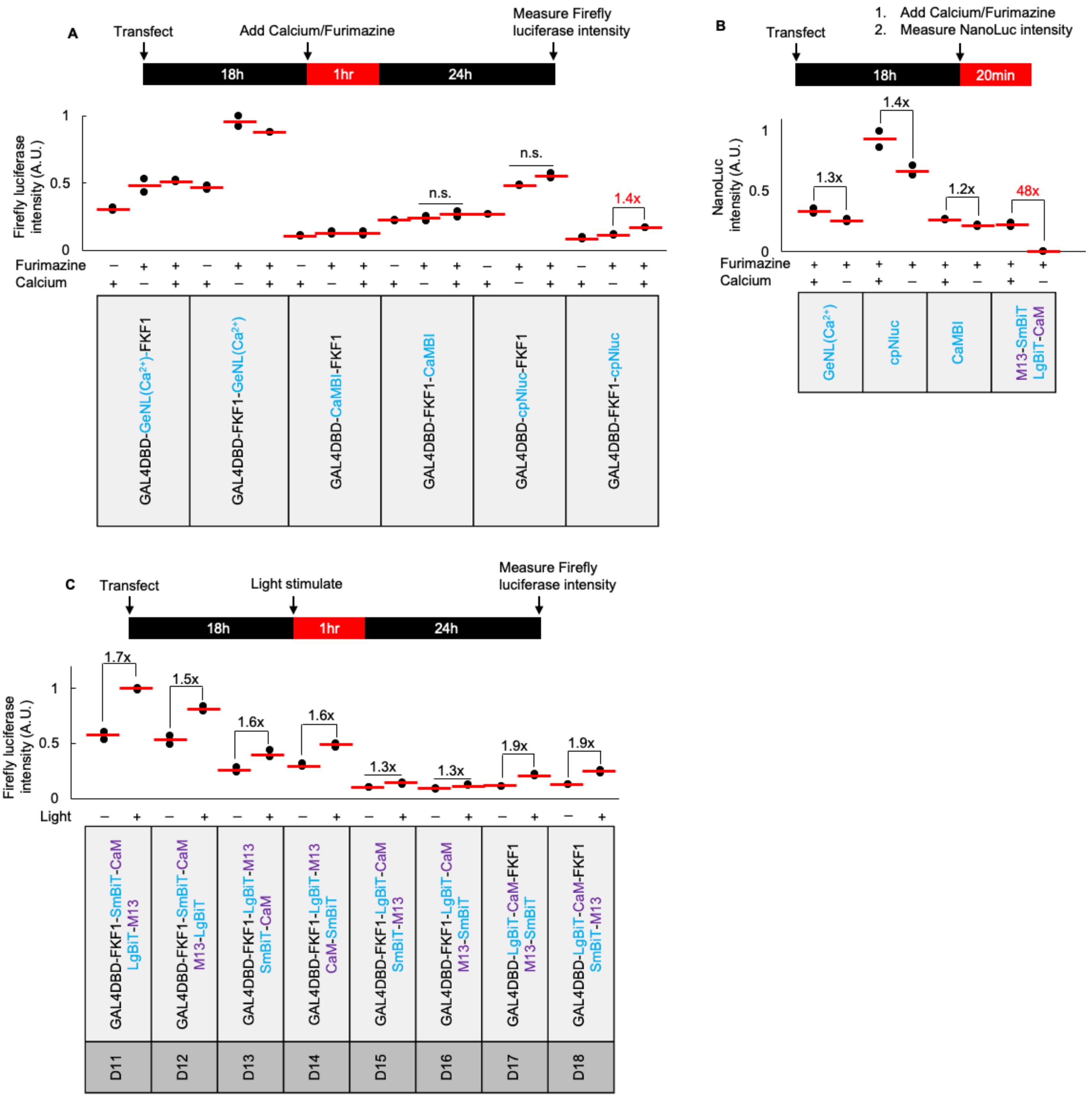
A
Transfect
Add Calcium/Furimazine
Measure Firefly luciferase intensity
18h
1hr
24h
Firefly luciferase intensity (A.U.)
1
0.5
0
n.s.
n.s.
1.4x
Furimazine
Calcium
GAL4DBD-GeNL(Ca²⁺)-FKF1
GAL4DBD-FKF1-GeNL(Ca²⁺)
GAL4DBD-CaMBI-FKF1
GAL4DBD-FKF1-CaMBI
GAL4DBD-cpNluc-FKF1
GAL4DBD-FKF1-cpNluc
B
Transfect
1. Add Calcium/Furimazine
2. Measure NanoLuc intensity
18h
20min
NanoLuc intensity (A.U.)
1.3x
1.4x
1.2x
48x
GeNL(Ca²⁺)
cpNluc
CaMBI
M13-SmBiT
LgBiT-CaM
C
Transfect
Light stimulate
Measure Firefly luciferase intensity
Firefly luciferase intensity (A.U.)
1.7x
1.5x
1.6x
1.6x
1.3x
1.3x
1.9x
1.9x
Light
GAL4DBD-FKF1-SmBiT-CaM
LgBiT-M13
GAL4DBD-FKF1-SmBiT-CaM
M13-LgBiT
GAL4DBD-FKF1-LgBiT-M13
SmBiT-CaM
GAL4DBD-FKF1-LgBiT-M13
CaM-SmBiT
GAL4DBD-FKF1-LgBiT-CaM
SmBiT-M13
GAL4DBD-FKF1-LgBiT-CaM
M13-SmBiT
GAL4DBD-LgBiT-CaM-FKF1
M13-SmBiT
GAL4DBD-LgBiT-CaM-FKF1
SmBiT-M13
D11
D12
D13
D14
D15
D16
D17
D18

**Supplementary figure 2 (related to figure 2). Testing single-component bioluminescent calcium sensors and alternative LuCID geometries A).** Alternative calcium-dependent NanoLuc designs tested in HEK293T cells with 1 hour furimazine +/- calcium. GeNL($Ca^{2+)}$ is calcium-enhanced NanoLantern (Suzuki et al., 2016). CaMBI is calcium-modulated bioluminescent indicator (Oh et al., 2019). cpNluc is circularly permuted NanoLuc with CaM-M13 inserted (Kim, personal communication). Two replicates per condition. **B**). Comparison of real-time bioluminescence intensities at their maximum (time = 4-6min mark after furimazine/calcium addition) from single-component calcium-dependent NanoLuc designs (GeNL($Ca^{2+}$), cpNluc, CaMBI) and split NanoLuc design. **C**). Remaining 8 construct combinations tested under 1 hour dark/light conditions from **Figure 2B**. FLuc reporter readout 24hrs later.

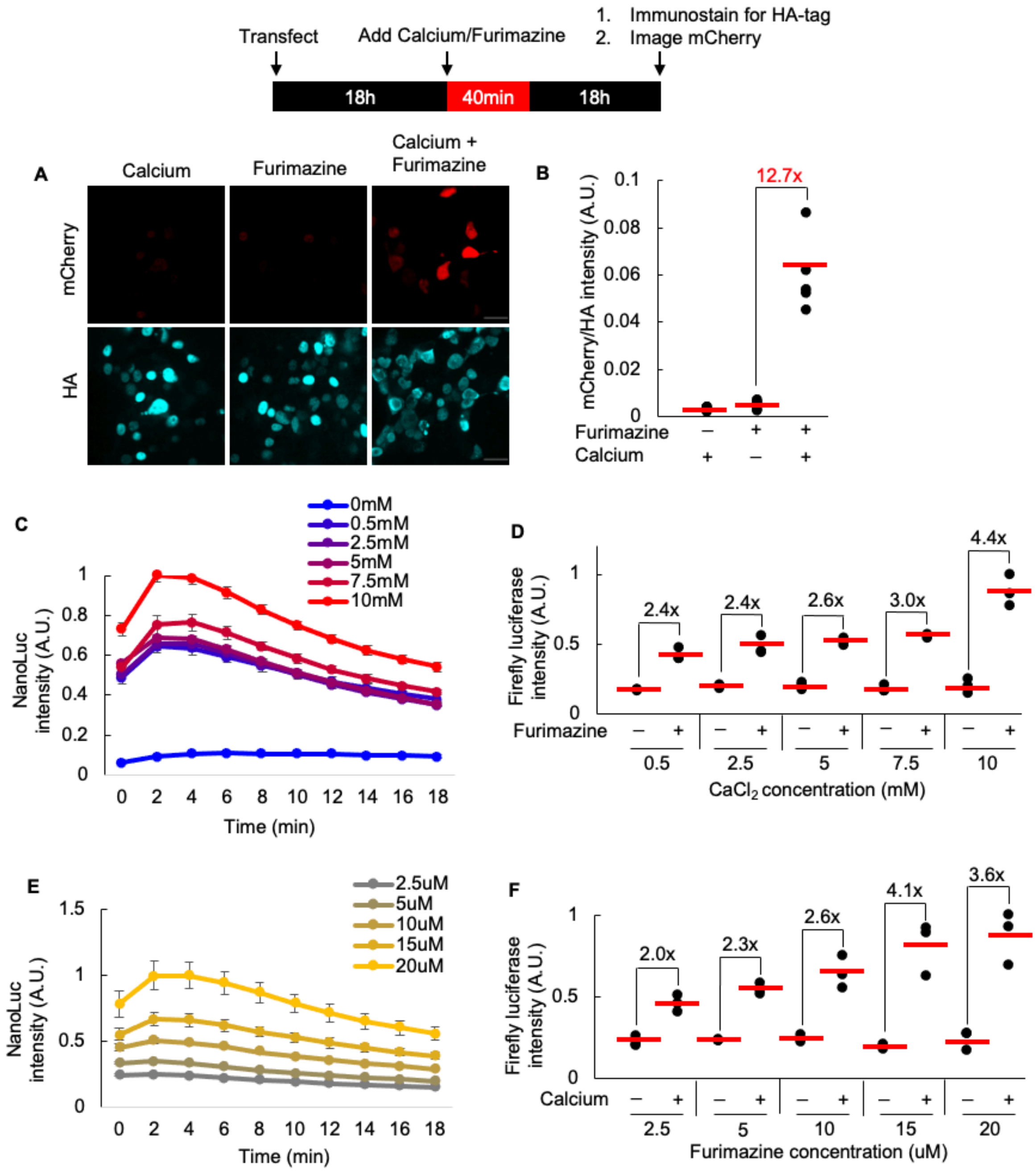

**Supplementary figure 3 (related to figure 3). Demonstrating alternative reporter and optimizing stimulation conditions A).** Example immunofluorescence images of mCherry expression in HEK293T cells 18 hours following furimazine treatment +/- calcium for 40 min (top, red). HA antibody staining labels GAL4DBD-M13-SmBIT-FKF1 (bottom, cyan). Scale bar = 30 μm**. B).** mCherry/HA fluorescence intensity ratios for all HA-positive HEK293T cells across 5 fields of view following 40 min of furimazine +/- calcium. Red line, mean. **C).** Titration of $CaCl_2$ concentration. Real-time NanoLuc intensity timecourse measurement in HEK293T cells expressing LuCID treated with furimazine and 2uM ionomycin +/- different $CaCl_2$ concentrations. **D).** FLuc reporter readout 24hours later under the same conditions as in (C). **E).** Titration of furimazine concentration. Real-time NanoLuc intensity timecourse measurement in HEK293T cells expressing LuCID treated with 2um ionomycin/5mM $CaCl_2$ + different furimazine concentrations. **F).** FLuc reporter readout 24hours later under the same conditions as in (E).

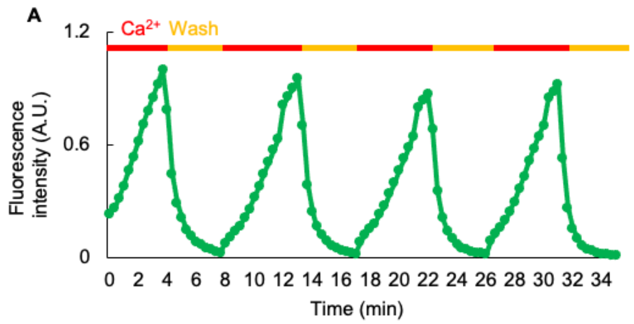

**Supplementary figure 4 (related to figure 4). Calcium measurement from HEK293T cells expressing GCaMP6s alone A**). Real-time measurement of calcium dynamics by GCaMP6s alone in HEK cells. Cells were stimulated repeatedly with 2uM ionomycin and 10mM $CaCl_2$ for 4 minutes and washed out for 2 min. Red bars above peaks indicate when ionomycin/$CaCl_2$ were added and yellow bars indicate when cells were washed out. This experiment was repeated twice.

**Supplementary Methods**

### Cloning

PCR fragments were amplified using Q5 polymerase (NEB, New England BioLabs). The vectors were double-digested with NEB restriction enzymes and ligated to gel-purified PCR products by Gibson assembly. XL1-Blue competent bacteria were heat shock transformed with the ligated plasmid products. All plasmids used in this paper are listed in **Table 1.** Full-length NanoLuc construct was a gift from Ute Hochgeschwender (Central Michigan University). The GI-VP16-IRES-GAL4DBD-FKF1 construct was a gift from Masuyaki Yazawa (Columbia University). The CaM and M13 genes were amplified from the GCaMP6s expression plasmid, obtained from Addgene (plasmid #100843). FKBP/FRB and GAI/GID were amplified from Addgene plasmids #40896 and #84240, respectively.

**Table 1.** Plasmids used in this paper

| Plasmid ID | Plasmid features | Plasmid vector | Promoter | Tags | Used in |
|---|---|---|---|---|---|
| p1 | NLS-GI-VP16-IRES-NLS-GAL4DBD-FKF1-NanoLuc | pcDNA3 | CMV | HA | Figure S1 |
| p2 | NLS-GI-VP16-IRES-NLS-GAL4DBD-NanoLuc-FKF1 | pcDNA3 | CMV | HA | Figures 1, S1 |
| p3 | NLS-GI-VP16-IRES-NLS-GAL4DBD-GeNL($Ca^{2+}$)-FKF1 | pcDNA3 | CMV | HA | Figure S2 |
| p4 | NLS-GI-VP16-IRES-NLS-GAL4DBD-FKF1-GeNL($Ca^{2+}$) | pcDNA3 | CMV | HA | Figure S2 |
| p5 | NLS-GI-VP16-IRES-NLS-GAL4DBD-CaMBI-FKF1 | pcDNA3 | CMV | V5/HA | Figure S2 |
| p6 | NLS-GI-VP16-IRES-NLS-GAL4DBD-FKF1-CaMBI | pcDNA3 | CMV | V5/HA | Figure S2 |
| p7 | NLS-GI-VP16-IRES-NLS-GAL4DBD-cpNanoLuc-FKF1 | pcDNA3 | CMV | HA | Figure S2 |
| p8 | NLS-GI-VP16-IRES-NLS-GAL4DBD-FKF1-cpNanoLuc | pcDNA3 | CMV | HA | Figure S2 |
| p9 | M13-SmBiT | pAAV | CMV | FLAG | Figure S2 |
| p10 | LgBiT-CaM | pAAV | CMV | FLAG | Figure S2 |
| p11 | NLS-GI-VP16-IRES-GAL4DBD-SmBiT-M13-FKF1 | pcDNA3 | CMV | V5/HA | Figure 2 |
| p12 | NLS-LgBiT-CaM | pAAV | CMV | FLAG | Figures 2, 3, 4, 5, S3 |
| p13 | NLS-CaM-LgBiT | pAAV | CMV | FLAG | Figure 2 |
| p14 | NLS-GI-VP16-IRES-GAL4DBD-CaM-SmBiT-FKF1 | pcDNA3 | CMV | V5/HA | Figure 2 |
| p15 | NLS-LgBiT-M13 | pAAV | CMV | FLAG | Figures 2, S2 |
| p16 | NLS-M13-LgBiT | pAAV | CMV | FLAG | Figures 2, S2 |

| | | | | | |
|---|---|---|---|---|---|
| p17 | NLS-GI-VP16-IRES-GAL4DBD-M13-LgBiT-FKF1 | pcDNA3 | CMV | V5/HA | Figure 2 |
| p18 | NLS-SmBiT-CaM | pAAV | CMV | FLAG | Figures 2, S2 |
| p19 | NLS-CaM-SmBiT | pAAV | CMV | FLAG | Figures 2, S2 |
| p20 | NLS-GI-VP16-IRES-GAL4DBD-CaM-LgBiT-FKF1 | pcDNA3 | CMV | V5/HA | Figure 2 |
| p21 | NLS-SmBiT-M13 | pAAV | CMV | FLAG | Figures 2, S2 |
| p22 | NLS-M13-SmBiT | pAAV | CMV | FLAG | Figures 2, S2 |
| p23 | NLS-GI-VP16-IRES-GAL4DBD-M13-SmBiT-FKF1 | pcDNA3 | CMV | V5/HA | Figures 2, 3, S3 |
| p24 | NLS-GI-VP16-IRES-GAL4DBD-FKF1-SmBiT-CaM | pcDNA3 | CMV | V5/HA | Figure S2 |
| p25 | NLS-GI-VP16-IRES-GAL4DBD-FKF1-LgBiT-M13 | pcDNA3 | CMV | V5/HA | Figure S2 |
| p26 | NLS-GI-VP16-IRES-GAL4DBD-FKF1-LgBiT-CaM | pcDNA3 | CMV | V5/HA | Figure S2 |
| p27 | NLS-GI-VP16-IRES-GAL4DBD-LgBiT-CaM-FKF1 | pcDNA3 | CMV | V5/HA | Figure S2 |
| p28 | NLS-GI-VP64-IRES-GAL4DBD-M13-SmBiT-FKF1 | pcDNA3 | CMV | V5/HA | Figure 3, 4, 5 |
| p29 | NLS-GI-VP16-IRES-GAL4DBD-FKBP-SmBiT-FKF1 | pcDNA3 | CMV | V5/HA | Figure 5 |
| p30 | NLS-LgBiT-FRB | pAAV | CMV | FLAG | Figure 5 |
| p31 | NLS-GI-VP16-IRES-GAL4DBD-GID-SmBiT-FKF1 | pcDNA3 | CMV | V5/HA | Figure 5 |
| p32 | NLS-LgBiT-GAI | pAAV | CMV | FLAG | Figure 5 |
| p33 | FLuc | pAAV | UAS | | Figures 1-5, S1-3 |
| p34 | mCherry | pAAV | UAS | | Figures S1, S3 |
| p35 | NLS-GCaMP6s | pAAV | CMV | FLAG | Figures 4, S4 |
| p36 | NLS-GI-VP16-IRES-GAL4DBD-FKF1 | pcDNA3 | CMV | HA | Figure S1 |
| p37 | NLS-NanoLuc | pAAV | CMV | HA | Figure S1 |
| Addgene #42500 | GI-GAL4DBD | pcDNA3 | CMV | | Figure S1 |
| Addgene #42499 | FKF1-VP16 | pcDNA3 | CMV | HA | Figure S1 |
| Addgene #135952 | NLS-CeNLuc | pEF1a-IRES- | EF1a | | Figure S1 |

| | | |
|---|---|---|
| | | DsRed-Express2 |

**HEK293T cell culture and transfection**

HEK 293T cells from ATCC were cultured as monolayers in Dulbecco's modified eagle medium (DMEM, Gibco) supplemented with 10% (vol/vol) fetal bovine serum (FBS, Sigma) and +1% (vol/vol) penicillin–streptomycin (PS) at 37 °C under 5% $CO_2$. Cells were grown in 24-well, 48-well or 96-well plates pretreated with 50 µg/mL human fibronectin (Millipore) for at least 10min. Cells were transfected at 60-90% confluence with 1mg/ml PEI max solution (polyethylenimine HCl max pH 7.3). Plasmid DNA was mixed with PEI max in serum-free DMEM and incubated at room temperature for 25-30min. Complete DMEM with 10% FBS + 1% PS was then added to the mixture and the entire volume was added to a well of a multi-well plate. Exact amounts of PEI, serum-free DMEM and complete DMEM used are summarized in **Table 2.** The plate was wrapped in aluminum foil and incubated in 37°C incubator for 18 hours.

**Table 2.** Plasmid amounts transfected in each experiment

| **Experiment type** | **Plasmid ID and amount per well** | **PEI (1mg/ml) amount per well (µL)** | **serum-free DMEM amount per well (µL)** | **Complete DMEM amount per well (µL)** | **Plate format** | **Used in** |
|---|---|---|---|---|---|---|
| BRET control of TF with full-length NanoLuc | 100ng p2 + 50ng p33 | 0.4 | 20 | 80 | 96-well | Figure 1d-e, S1a |
| BRET control of TF with full-length NanoLuc | 100ng p1 + 50ng p33 | 0.4 | 20 | 80 | 96-well | Figure S1a |
| BRET control of TF with full-length NanoLuc + mCherry reporter | 200ng p2 + 100ng p34 | 0.7 | 30 | 170 | 48-well | Figure S1b-c |
| BRET proximity: NanoLuc fused to FKF1 | 500ng p2 + 250 ng p33 | 1.8 | 70 | 330 | 24-well | Figure S1e |
| BRET proximity: NanoLuc unfused to FKF1 | 500ng p36 + 500ng p37 + 250ng p33 | 3 | 120 | 280 | 24-well | Figure S1e |

| | | | | | | |
|---|---|---|---|---|---|---|
| BEACON activity + localization | 500ng Addgene #135952 + 500ng Addgene #42500 + 250ng Addgene #42499 + 250 ng p33 | 3.6 | 140 | 260 | 24-well | Figure S1e-g |
| GAL4DBD-FKF1 localization | 500ng p36 | 1.2 | 50 | 350 | 24-well | Figure S1g |
| Unfused NanoLuc localization | 500ng p37 | 1.2 | 50 | 350 | 24-well | Figure S1g |
| Single-chain bioluminescent calcium indicators | 100ng of p3/p4/p5/p6/p7/p8 + 50ng p33 | 0.4 | 20 | 80 | 96-well | Figure S2a-b |
| Split-NanoLuc bioluminescent calcium indicator | 100ng p9 + 100ng p10 | 0.5 | 20 | 80 | 96-well | Figure S2b |
| LuCID construction | 100ng of each indicated component in figure 2 and S2c + 50ng p33 | 0.6 | 20 | 80 | 96-well | Figure 2, S2c |
| LuCID stimulation and expression condition optimization | 100ng p23 + 100ng p12 + 50ng p33 | 0.6 | 20 | 80 | 96-well | Figure 3a-b, S3c-f |
| LuCID plasmid ratio optimization | 1. 200ng p23 + 200ng p12 + 100ng p33 (1:1:0.5)<br>2. 200ng p23 + 100ng p12 + 100ng p33 (1:0.5:0.5)<br>3. 100ng p23, p12, p33 (0.5:0.5:0.5)<br>4. 200ng p23 + 200ng p12 + 10ng p33 (1:1:0.05) | 1. 1.2<br>2. 1<br>3. 0.7<br>4. 1<br>5. 1.2<br>6. 1.2 | 1. 50<br>2. 40<br>3. 30<br>4. 40<br>5. 40<br>6. 40 | 1. 150<br>2. 160<br>3. 170<br>4. 160<br>5. 160<br>6.160 | 48-well | Figure 3c |

| | | | | | | |
|---|---|---|---|---|---|---|
| | 5. 200ng p23 + 200ng p12 + 5ng p33 (1:1:0.025)<br>6. 200ng p28 + 200ng p12 + 5ng p33 | | | | | |
| LuCID with mCherry reporter | 200ng p28 + 200ng p12 + 5ng p34 | 1.2 | 40 | 160 | 48-well | Figure S3a-b |
| LuCID comparison to GCaMP6s | 100ng p28 + 100ng p12 + 100ng p35 or 100ng p35 alone | 0.7 or 0.3 | 30 or 10 | 70 or 90 | 96-well | Figure 4a, S4 |
| LuCID characterization | 100ng p28 + 100ng p12 + 2.5ng p33 | 0.5 | 20 | 80 | 96-well | Figure 4b-c |
| LuCID with SERCA inhibitors and activator | 100ng p28 + 100ng p12 + 2.5ng p33 | 0.5 | 20 | 80 | 96-well | Figure 5a-b |
| LuCID with FKBP/FRB system | 100ng p29 + 100ng p30 + 2.5ng p33 | 0.5 | 20 | 80 | 96-well | Figure 5d-e |
| LuCID with GAI/GID system | 100ng p31 + 100ng p32 + 2.5ng p33 | 0.5 | 20 | 80 | 96-well | Figure 5f-g |

**HEK293T fixation and immunostaining**

After transfection, stimulation and incubation for 18 hours for mCherry reporter expression (as indicated above), cells were fixed in 4% paraformaldehyde (PFA) in PBS for 15 min at room temperature. Cells were permeabilized by incubation with cold methanol at −20 °C for 7 min, washed three times with room-temperature PBS, then immunostained with rabbit-anti-HA antibody to detect FKF1-containing components (1:1,000 dilution, Rockland) in 2% BSA solution in PBS for 45 min at room temperature with gentle rocking. The cells were washed twice with PBS and then incubated with a secondary antibody, anti-rabbit-Alexa Fluor 647 (1:1,000 dilution, Life Technology) in 2% BSA solution in PBS, for 20 min at room temperature. Cells were washed twice with PBS and maintained in PBS at 4 °C until imaging.

**Fluorescence microscopy of HEK293T cells**

Confocal imaging was performed on a Zeiss AxioObserver inverted confocal microscope with 40x and 63x oil-immersion objectives, outfitted with a Yokogawa spinning disk confocal head, a Quad-band notch dichroic mirror (405/488/568/647), and 405 (diode), 491 (DPSS), 561 (DPSS) and 640-nm (diode) lasers (all 50 mW). The following combinations of laser excitation and emission filters were used for the fluorophores: mCerulean3 (405 nm laser excitation, 445/40 nm emission), Alexa-Fluor-647 (647 nm laser excitation, 680/30 nm emission) and mCherry (561 laser excitation; 617/73 emission). Acquisition time was 100ms. All images were collected and processed using SlideBook (Intelligent Imaging Innovations). 5 fields of view were acquired per condition. A mask was defined using anti-HA immunofluorescence (expression of the FKF1 component). Mean mCherry intensity was calculated within this mask. A second mask outside of HA immunofluorescence was drawn to define background mCherry mean intensity. Background mCherry intensity outside of HA immunofluorescence was subtracted from mCherry intensity within the HA mask. Background-corrected mCherry intensity normalized by HA immunofluorescence intensity across 5 fields of view are reported for each condition.
